# Graph representations of iEEG data for seizure detection with graph neural networks

**DOI:** 10.1101/2023.06.02.543277

**Authors:** Alan A. Díaz-Montiel, Milad Lankarany

## Abstract

Epilepsy is a neurological disorder that affects over 50 million individuals worldwide. Today, the gold-standard treatment for those who are drug resistant, meaning that symptoms cannot be controlled with medication, is to surgically remove the seizure onset zone (SOZ), the area of the brain believed to cause seizures: the main symptom of epilepsy. Unfortunately, around 50% of drug resistant patients are not resective candidates, which can be attributed in part to poor SOZ localization. SOZ localization is a complex and lengthy procedure, requiring visual inspection and manual processing by human experts that first need to localize and isolate seizure events. The intracranial electroencephalography (iEEG) is a tool that records electrophysiological activity of the inner brain at different regions and depths, and provides critical information on the SOZ. However, iEEG data processing methodologies are not standardized, and practice and resources vary across hospitals and clinics. To assist human experts with systematic processing of iEEG data, we propose a data processing pipeline that generates graph representations of iEEG data. We evaluate 9 different graph representations of publicly available iEEG data from 25 patients with epilepsy with a graph neural network model trained to detect seizures. Our results suggest that graph representations of iEEG data that leverage electrode and functional connectivity features are powerful data structures to analyze and interpret iEEG data in the context of epilepsy. We anticipate that our data pipeline that provides a systematic processing of neural data with graphs can integrate other data modalities like neuroimaging data. Moreover, methods used in the data pipeline have potentials to apply to other neurological disorders such as Parkinson’s disease or major depression disorder.

## 1 Introduction

Epilepsy is an incurable neurological disorder that affects over 50 million individuals worldwide [1]. While symptoms can often be controlled with prescribed medication and provide good quality of life to patients, around 30% of people with epilepsy are medically refractory, thus their symptoms cannot be controlled with medication [2]. For this population, the gold-standard treatment is resective surgery, a procedure that consists in removing a brain region that is believed to cause seizures: the main symptom of epilepsy [3]. This brain region is referred to as the seizure onset zone (SOZ), and its localization is a complex and lengthy task performed by expert epileptologists. Unfortunately, resective surgery success rate is about 50-70%, which can be partly attributed to poor SOZ localization[2].

Patients with epilepsy are assessed within the Epilepsy Monitoring Unit (EMU) of a hospital where they undergo 24/7 video and electroencephalographic (EEG) monitoring for up to 4 weeks, although in rare occasions patients may undergo monitoring periods of up to 3 months. Epilepsy is a clinical-electroencephalographic diagnosis [4], with the EEG providing critical information for the localization of the SOZ. These recordings generate a large amount of video and EEG recordings. A portion of these patients will also undergo intracranial EEG (iEEG) monitoring where electrodes are implanted within the brain to determine where seizures are arising from [3]. Annually, around 300 patients undergo EEG recordings, and close to 40 patients undergo iEEG recordings at the EMU of the Toronto Western Hospital in Toronto, ON, Canada.

Crucial steps for the localization of the SOZ from iEEG recordings are signal processing and data management procedures. For instance, to identify epileptogenic zones, the brain areas with high concentration of seizure activity, it is necessary to accurately detect seizure activity within the data. Due to the sophisticated surgery for electrode implantation, the limited number of electrodes for monitoring neural activities, and the current manual processing clinical methodologies, the need for improving the reliability of seizure detection and SOZ localization in an automatic way is urgent and timely. For this, several tools have been proposed, including heuristic and machine learning algorithms [5–8]. However, these methods mainly suffer from the lack of scalability and interpretability that limit their usability for clinical translation and practices [5,6]. Also, the vast amount of works found in the literature mainly used scalp EEG recordings, which is in part due to a lack of openly available, de-identified datasets of iEEG recordings.

In recent years, graph neural networks (GNNs), a paradigm of artificial neural networks optimized to operate on graph-structured data, have shown great potential as a tool to study how the brain represents information and produces cognition and behavior [9–12]. In this paper, we investigate the use of GNNs to automate the clinical process of seizure detection.

GNNs operate similarly to convolutional neural networks (CNNs), with the key difference being the structure of the input data from which these models *learn*. While CNNs operate on grid-structured data, for example, matrices and images, GNNs operate on graph-structured data contained within a **graph representation (GR)**. A *GR* = (*A*, **V, E**), is a data structure composed of 3 elements: an adjacency matrix, *A*^*N×N*^ ; a node features vector, **V** = ℝ^*N×D*^; and an edge features vector, **E** = ℝ^*N×N×D*^*′*, where *N* represents the nodes in the graph, *D* and *D*^*′*^ correspond to the number of features considered for each features vector. Consequently, to understand the full potential of GNNs for seizure detection and SOZ localization, it is utterly important to clearly define how to use iEEG data to increase the representational power of graphs. In this study, we aim to provide answers to this by evaluating multiple iEEG-GRs with different node and edge features from the iEEG data, and assess their impact to seizure detection with GNNs.

We use a recently validated GNN model from [8] to benchmark the performance of our proposed iEEG-GRs. In [8], Grattarola et. al. proposed a 2-layer binary classification GNN model with an edge-conditioned convolution layer (ECC)[13] followed by a graph attention layer (GAT)[14]. In a nutshell, the model determines whether an iEEG-GR represents ictal activity (seizure, class 0) vs. non-ictal activity (non-seizure, class 1). To create the iEEG-GRs, the authors used functional connectivity networks (FCNs) of iEEG data. FCNs are network abstractions of brain data that assess the degree of “connectivity” across brain regions by comparing the occurrence of simultaneous activity over periods of time. From iEEG data, FCNs are computed by comparing the dynamics across signals, where higher synchronicity in the activity is interpreted as higher connectivity. For instance, FCNs in [8] were created using the Pearson correlation and the phase-lock value (PLV) methods, which assess the similarities in energy levels and signal oscillatory patterns across signals, respectively. However, the authors did not use node and edge feature vectors to leverage the iEEG data, and they set these as all-ones vectors: *GR*^*′*^ = (*A*, **1, 1**). They reported an average seizure detection precision-recall of 81.24 ± 10.37 across 8 patient records. We hypothesize that by leveraging iEEG signal data and functional connectivity measurements as node and edge feature vectors of GRs, we can improve the seizure detection performance of the GNN model.

In this study, we propose a data management and processing pipeline to generate iEEG-GRs. We have integrated the GNN model proposed in [8] within our pipeline, and use it to evaluate 9 iEEG-GR and quantified their impact for seizure detection as a binary classification problem. Moreover, we evaluate the 9 GRs and their impact to detect preictal (before seizure), ictal (during seizure), and postictal (after seizure) activity as a multi-class classification problem. We used a publicly available dataset of patients with epilepsy hosted at OpenNeuro, which consists in iEEG seizure recordings from 25 patients with epilepsy collected across 4 epilepsy centers in the US [15]. We show that by leveraging energy-at-electrode information as node features and functional connectivity measurements as edge features, we can improve the performance of the GNN model to detect seizure activity as a binary classification problem with an average accuracy of 91%, an area under the curve (AUC) of 95% and an F1-score of 90%. For the multi-class classification problem, we obtained an average accuracy of 88%, an AUC of 96%, and an F1-score of 86%. Our concrete contributions are summarized as follows:

- We propose a novel data processing pipeline to generate GRs of iEEG data.
- We integrate the GNN model proposed in [8] for seizure detection within our pipeline, and extend it to account for node and edge features pertinent to the iEEG electrodes and functional connectivity measurements.
- We extend the evaluation of the GNN model in [8] by testing it with a publicly available dataset of patients with epilepsy[15].
- We identify and quantify iEEG data patterns across 25 epilepsy patients and their impact to build GNN models for seizure detection.
- We build a GNN model for detecting preictal, ictal, and postictal iEEG signal activity.
- We assess what graph representation reaches the best accuracy, AUC, and F1-score for seizure detection.

## 2 Methods and materials

### 2.1 On the data

We used a publicly available dataset hosted at OpenNeuro with Accession Number ds003029 [15], which consists in iEEG and EEG data from 100 individuals across 5 epilepsy centers in the US that underwent resective surgery. However, due to reasons that we disclose in the Discussion section, for this study we only use the data from 25 patients across 4 epilepsy centers. This dataset was first used in [7], where Li et. al. proposed the metric of neural fragility as a biomarker of epilepsy that can be used for SOZ localization. In Table 1 we show a summary of the information related to the extract of the dataset that we used, which contains demographic information about the patients such as race, age, and gender, as well as information on the epilepsy cases such as ILAE score, Engel score, and surgery outcome. The surgery outcome was set as success or failure, where the former meant that no seizure events were registered on the patient after surgery, and the latter that seizure events continued to occur. The post-surgery monitoring period varied from patient to patient, with the shorter monitoring period being 1 year and the longest 7 years.

**Table 1:** Summary of 25 patient information from the OpenNeuro ds003029 dataset [15].

| Race | Mean age at surgery | Gender | ILAE score | Engel score | Surgery outcome |
| --- | --- | --- | --- | --- | --- |
| 6 caucasian | $36.4 \pm 9.9$ | 8 females | 12, score 1 | | |
| 3 african american |  | 13 males | 3, score 2 | 14, score 1 | 14 success |
| 3 hispanic | 4 patients NA | 4 NA | 1, score 3 | 3, score 2 | 8 failure |
| 13 NA |  |  | 2, score 4 | 1, score 3 | 3 NA |
|  |  |  | 1, score 5 | 4, score 4 |  |
|  |  |  | 3, score 6 | 3, NA |  |
|  |  |  | 3, NA |  |  |

Each patient dataset consists in 1 to 4 runs of seizure activity of varying duration. A run is considered as a single recording of iEEG activity that captures one seizure event, and is characterized by containing three time periods, the preictal, ictal, and postictal periods, which correspond to instances of time before, during, and after the seizure event. Each run that we selected has been clinically annotated for seizure onset and seizure offset times. The monitoring resolution across patients varied from 250 Hz, 500 Hz, and 1 kHz. An example of a run of one patient is illustrated in Figure 1, where the y-axis represents the iEEG electrodes at different brain regions, and the x-axis represents time in seconds. We have 86 runs in total for the 25 patients, with 3 ± 1 runs in average per patient. The average number of electrodes across the 25 patients is 115 ± 22, and the average run duration time in seconds is 264 ± 170. The preictal, ictal, and postictal period duration also varies across runs.

**Figure 1.**
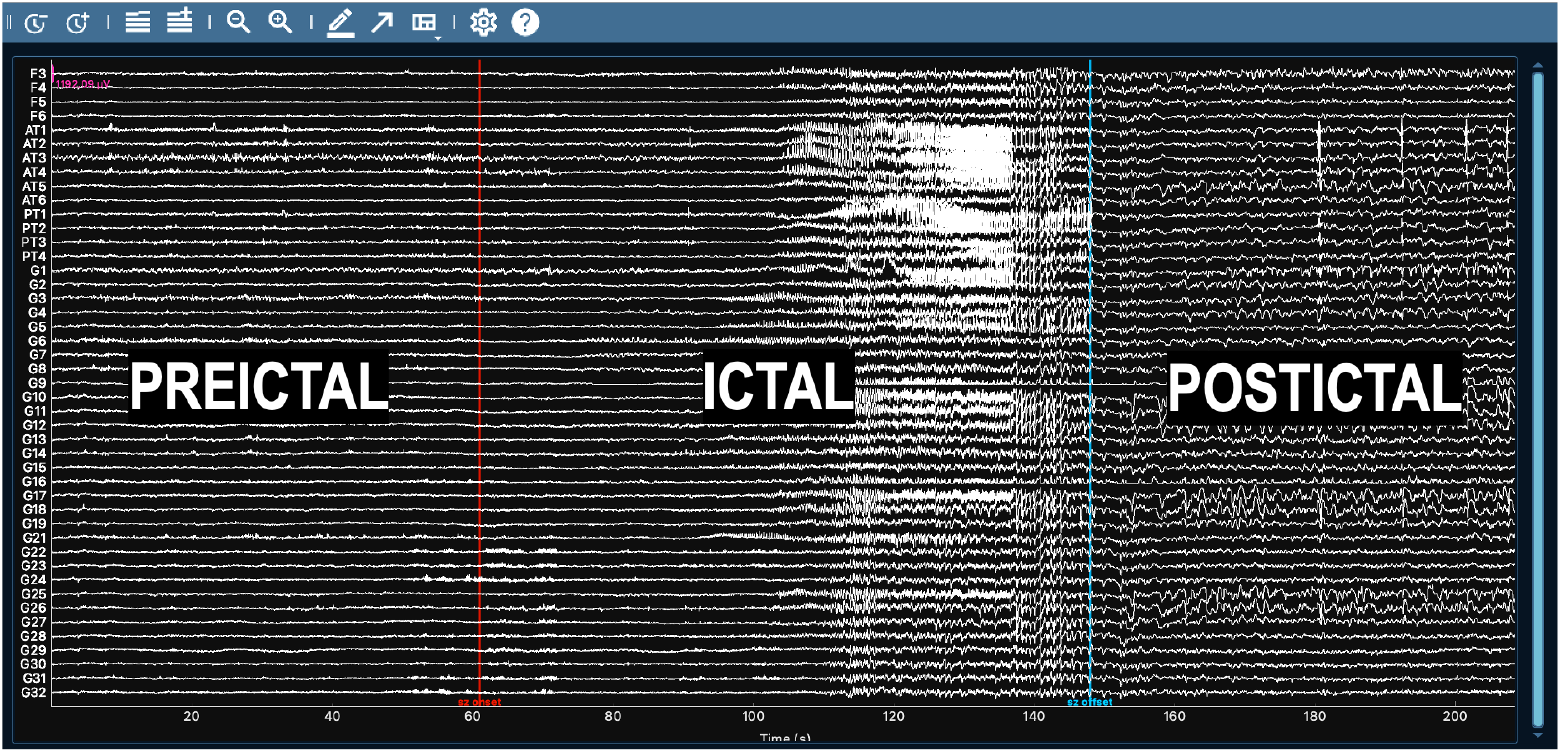
Example of an iEEG run from one patient illustrating the preictal, ictal, and postictal signal periods, divided by the seizure onset and seizure offset clinical markers in red and blue, respectively. Figure generated with MNE [16].

### 2.2 Data processing pipeline

The data processing pipeline that we developed is illustrated in Figure 2. It is entirely developed in Python, and it uses the MNE software [17] for iEEG data management and preprocessing, the Spektral project [18] and Keras [19] with Tensorflow [20] for data balancing and GNN model handling, and it leverages the supercomputer infrastructure from the Canada-wide High Performance Computing platform from the Digital Research Alliance of Canada. The technical integration of these tools with our customized software interfaces will be disseminated elsewhere.

**Figure 2.**
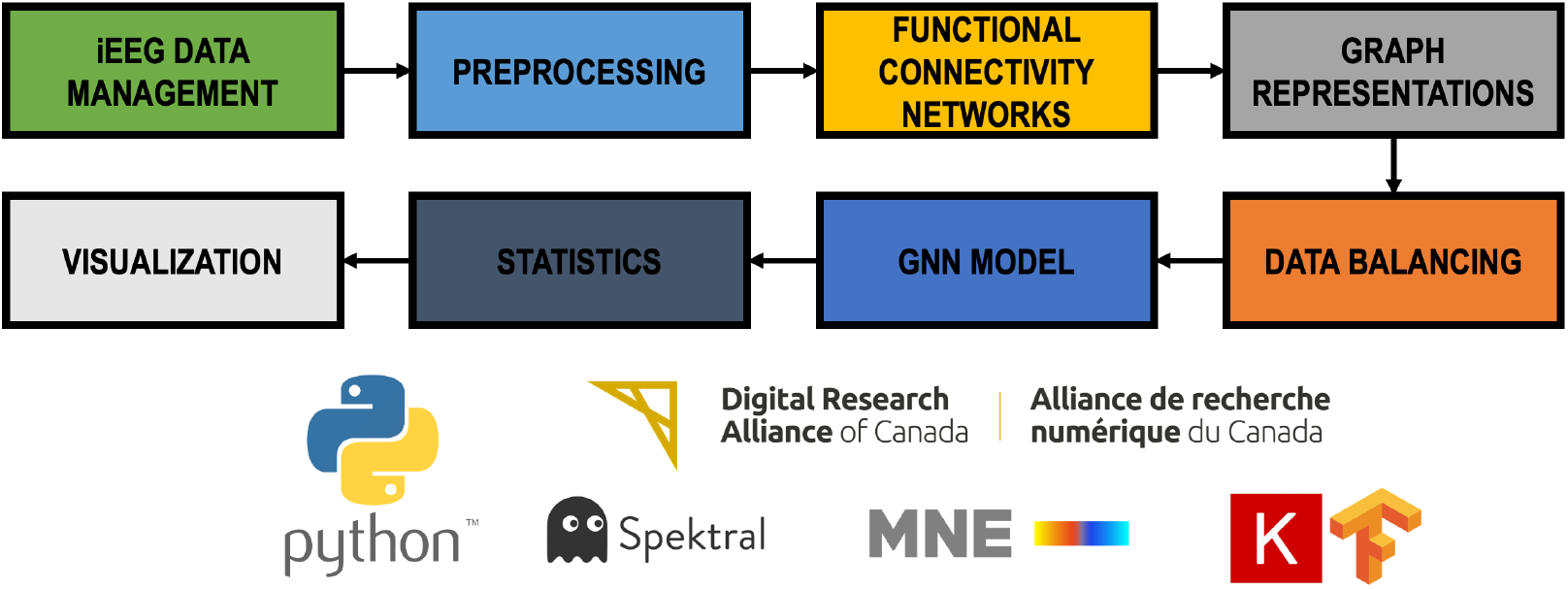
Diagram illustrating our data pipeline. The tool is entirely developed in Python, and it is powered by MNE [17] for iEEG data management and preprocessing, Spektral [18] and Keras [19] with Tensorflow [20] for data balancing and GNN model handling, and it is running on the Canada wide High Performance Computing platform managed by the Digital Research Alliance of Canada.

#### 2.2.1 iEEG data management

Monitoring an epilepsy patient with iEEG for a 24/7 period collects around 10 TB (terabytes) of data. Handling iEEG data is a concern of every hospital or EMU, and it is influenced by the monitoring equipment available to these entities, and the clinical and technical expertise of the individuals working with the data. This unstandardized practice has resulted in large heterogeneous iEEG datasets that are not Findable, Accessible, Interoperable and Reusable (FAIR), and that have limited clinical use within their own centers. Thus, iEEG data management standardization protocols are required. In 2019, the iEEG-BIDS protocol was proposed in [21], which extends the Brain Imaging Data Structure (BIDS) protocol [22] to operate with iEEG data. Since then, the iEEG-BIDS protocol has been spread around the iEEG research and clinical community, although its adoption as a gold-standard has yet to be realized. The OpenNeuro ds003029 dataset [15] is stored under the iEEG-BIDS format. Within our pipeline, we leverage iEEG-BIDS data handling processes with the MNE software [16], which provides robust tools to read/write and process this type of data.

#### 2.2.2 Data preprocessing

Although iEEG data can be captured at high sampling rates of up to 25KHz [23], clinical practice often relies on lower recording resolutions from 250Hz to 2kHz [24]. Despite the availability of high resolution sensors, the monitoring process is far from perfect, and recordings are often corrupted due to many-form noise sources. These include power-line electricity noise and artifacts caused by involuntary body movements (i.e., eye-lid or muscle movement). Moreover, it is common to have unusable data from bad channels due to bad electrode placement or contact. Data corruption phenomena are commonly fixed by preprocessing the data, which is yet another unstandardized process in clinical and research practice.

Our iEEG data preprocessing pipeline consists in first removing bad channels from the dataset which were identified by the clinicians, and are part of the metadata found within the iEEG-BIDS format. Next we apply a notch filter at 60 Hz and corresponding harmonics (120 Hz, 180 Hz, etc.), to attenuate the presence of power-line noise. We do not perform any artifact removal or artifact reconstruction method as we are interested in using the data with as little preprocessing as possible. This part of our pipeline also leverages the MNE software tools for iEEG signal processing.

The electrophysiological activity of the brain is known to use different frequency bands for different purposes. For instance, recently low-gamma oscillations have been found to be involved in emotional regulation, and high-gamma oscillations seem to be involved in speech temporal information [25,26]. However, the self-regulatory neuromodulation processes of the brain are vastly unknown. Consequently, we are also interested in investigating the role of different frequency bands and their role in seizure events. Moreover, we are interested in devising how to use this information to create powerful iEEG-GRs to improve seizure detection with GNNs. For this, our preprocessing pipeline also includes bandpass filtering with a highpass filter set at 0.1 Hz and a lowpass filter set at the Nyquist limit, which depends on the sampling frequency of each run. Last, we clip each preprocessed run in their preictal, ictal, and postictal signal traces, and stored them in serialized file objects.

#### 2.2.3 Functional connectivity networks

FCNs are abstractions of brain data that aim to represent the dynamics of neurophysiological activity recorded with neuroimaging tools, such as diffusion tension imaging (DTI) or electrophysiological tools, such as iEEG. FCNs are useful to study neurophysiological activity from a networks perspective, and they can be used to map network-modeled neurophysiological activity to behavioral and cognitive dimensions. Network analysis of iEEG data pose new perspectives to uncover neural circuits underlying neurological disorders, which will be instrumental for the development of new treatment options[27].

In this study, we focus solely on iEEG-based FCNs. Thus, the purpose of the iEEG data-to-network abstraction is to quantify the degree of similarity across iEEG signals. While there are several methods to create iEEG-based FCNs, mapping FCNs to behavioral or cognitive tasks is an active research field, and there is no one method that is useful for all cases. For this study, we consider the methods of Pearson correlation, which is used to measure the similarity of energy levels across signals over time, in the time domain; the coherence, which also measures similarity of energy levels across signals over time but in the frequency domain; and the phase-lock value (PLV), which measures where are the signals over time.

Our method for FCN creation is illustrated in Figure 3. In block **A**, we show an iEEG run from one patient, where the y-axis depicts the signals collected at each electrode (measuring energy in Volts), and the x-axis depicts time starting at *t*0 and ending at *L* seconds. For the binary classification problem we label nonictal data (preictal and postictal traces) as class 0, and ictal data as class 1. For the multi-class classification problem we label preictal, ictal, and postictal data as class 0, 1, and 2, respectively. There are two markers on each run, *t*_*on* and *t*_*off*, indicating the beginning and the end of the ictal activity as annotated by the clinical experts. To compute FCNs, we define a window, *W*, depicted in green at the top-left, which indicates the interval of time in which the degree of connectivity across signals is assessed. Then, we define a sliding window, *SW*, which indicates how to slide *W* across the run to create FCN sequences as depicted in blocks **B** and **C**. For this study, we consider 1 second *W* and 0.125 seconds *SW*.

**Figure 3.**
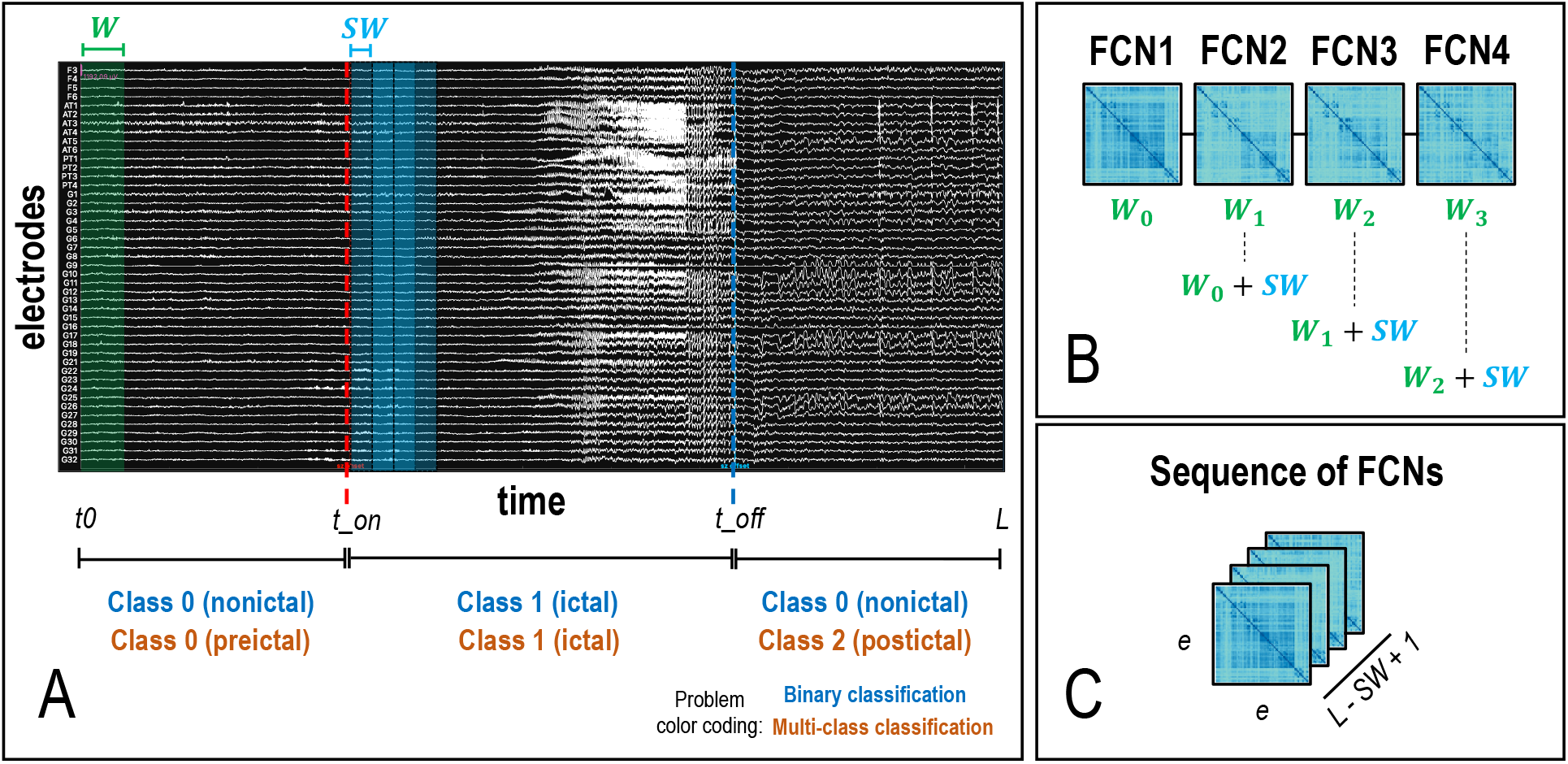
Methodology to create functional connectivity networks (FCNs). **A**. Every iEEG record (each run) of duration *L* has 2 main time marks, the seizure onset (*t*_*on*) and the seizure offset (*t*_*off*). The signal trace within *t*0 and *t*_*on* indicates the preictal period of the signal, which we label as 0 for the binary and multi-class classification problems. The signal trace within *t*_*on* and *t*_*off* indicates the ictal period, which we label as 1 for the binary and multi-class classification problems. The signal trace within *t*_*off* and *L* indicates the postictal period, which we label as 0 and 2 for the binary and multi-class classification problems, respectively. To create FCNs, we declare a window *W* (in green, top-left) that indicates the portion of the iEEG record to analyze. Then, we declare a sliding window *SW* (in blue, top-center) that indicates how to slide *W* over the entire record. **B**. Illustration of 4 FCNs sequentially created by sliding *W* by *SW*. **C**. The result is a multidimensional array shaped by (*e, e, L − SW* + 1), representing an FCN sequence.

#### 2.2.4 Graph representations

In computer science, GRs are data structures that extend the original graph data structure to account for node and edge feature vectors. Take for instance a regular graph data structure illustrated in Figure 4, **A**, where *G* = *{V, E}*, and *V* and *E* are sets of vertices and edges between vertices. Instead, the *GR* in **B** is represented as 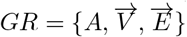, where *A* is an adjacency matrix or “original graph”, 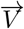 and 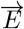 are sets of multidimensional vectors representing node and edge features [28]. As shown in block **C**, similarly to the creation of sequences of FCNs, we can create sequences of GRs to increment the representational power of the abstracted data.

**Figure 4.**
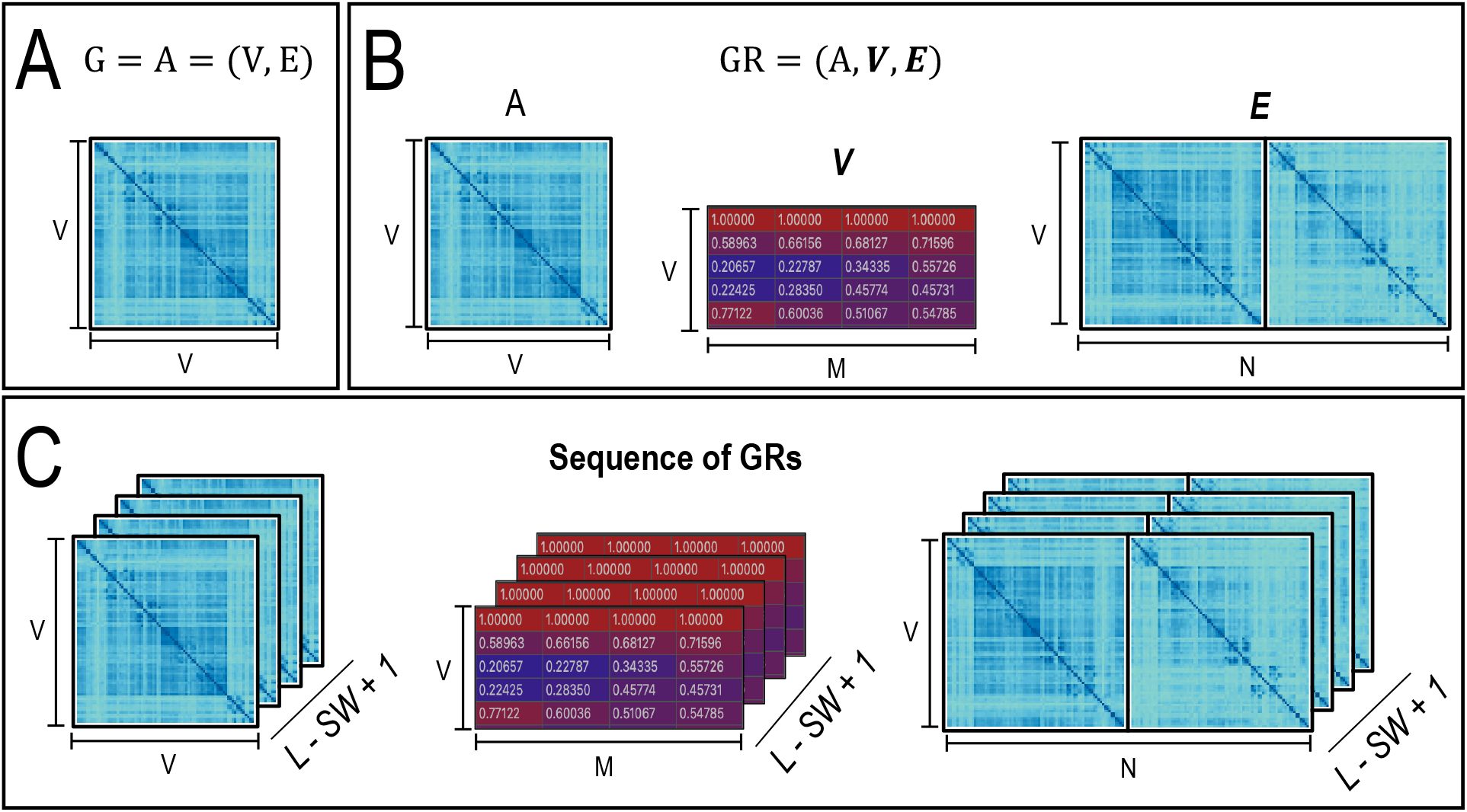
Illustration of graph representations (GRs). **A**. *G* = *A* = (*V, E*) is the original graph represented as a set of vertices *V* and edges *E*. **B**. *GR* = (*A*, **V, E**) is a graph representation with 3 elements: and adjacency matrix, *A*^*V ×V*^, a node features vector, *V* ^*V ×M*^, and an edge features vector, *E*^*V ×N*^, where *M* and *N* are the number of node and edge features, respectively. **C**. Similar to the FCN sequences, we can create sequences of GRs.

Graph-structured data contained within GRs are commonly used as input data to build GNN models, a paradigm of artificial neural networks optimized to operate with graph-structured data. In recent years, GNN models have been used to advance the understanding of real-life network phenomena found across several scientific fields including physics and chemistry [28]. More recently, some works have started to show the potential of GNNs to assist the field of neuroscience [8,29], but further research is require to deploy these models in the clinical practice. Importantly, the ability of GNN models to learn about the graphs they operate on is dependant on the abstraction of the data within the GRs, which is still considered an art in the realm of deep learning and AI. In this study, we investigate the creation of GRs of iEEG data that are more helpful to build a GNN model for seizure detection.

Our goal is to understand what combination of graph representation elements renders the most powerful iEEG-GR for seizure detection. We consider 9 GRs, presented in Table 2, which we cluster in 3 groups. Group 1 is composed of tests b1, b2, and b3 correspond to the baseline GRs used in [8] (b1 and b2), where node and edge features are considered as all-ones vectors, and adjacency matrices are considered as FCNs of the Pearson correlation and PLV methods. Test b3 was not used in [8], but we include it to evaluate the usage of the coherence method in this form. Group 2 is composed of tests t11, t12, and t13, and correspond to GRs that use the average energy recorded by the iEEG electrodes as node features vector and FCNs of the Pearson correlation, coherence, and PLV methods. The average energy at electrode vector has a shape of (*V*, 1), where V is the number of electrodes. Last, group 3 are tests t21, t22, and t23 and correspond to GRs that consider the average energy at electrode and the average energy at electrode by frequency band as node features vector. The average energy at electrode by frequency band has a shape of (*V, F*), where *F* corresponds to the frequency bands considered. For iEEG signals that were recorded with a 250 Hz sampling rate, *F* = 6, as we consider the frequency bands *δ*, delta (1 - 4 Hz), *θ*, theta (4 - 8 Hz), *α* (8 - 13 Hz), alpha *β*, (13 - 30 Hz), *γ*, gamma (30 - 70 Hz), and Γ, high-gamma (70 - 100 Hz). For signals that were recorded with a 500 Hz sampling rate, *F* = 7, as we also consider ripples in the frequency band (100 - 250 Hz). For signals recorded at 1 kHz sampling rate, *F* = 8, as we also consider fast ripples in the frequency band (250 - 500 Hz). We then create the node features vectors for these tests by concatenating the vectors for the average energy at electrode and the average energy at electrode by frequency band. Similarly, for the edge features vector we concatenate 2 FCNs that are different from the adjacency matrix. For example, in test t21, the adjacency matrix is considered to be a Pearson correlation-based FCN, while the edge features vector is the concatenation of the coherence FCN with the PLV FCN.

**Table 2:** Graph representation elements.

| Test | Adjacency matrix | Node features | Edge features |
| --- | --- | --- | --- |
| b1 | Pearson correlation | All-ones vector | All-ones vector |
| b2 | Phase-lock value | All-ones vector | All-ones vector |
| b3 | Coherence | All-ones vector | All-ones vector |
| t11 | Phase-lock value | Average energy at electrode | Pearson correlation |
| t12 | Pearson correlation | Average energy at electrode | Phase-lock value |
| t13 | Phase-lock value | Average energy at electrode | Coherence |
| t21 | Pearson correlation | Average energy at electrode<br>Average energy at electrode by frequency band | Coherence<br>Phase-lock value |
| t22 | Phase-lock value | Average energy at electrode<br>Average energy at electrode by frequency band | Pearson correlation<br>Coherence |
| t33 | Coherence | Average energy at electrode<br>Average energy at electrode by frequency band | Pearson correlation<br>Phase-lock value |

#### 2.2.5 Data balancing

To evaluate the GNN model, we split each patient dataset in train, validation, and test sets, taken from 80%, 10%, 10% of the data, respectively. However, the amount of data samples varies from the preictal, ictal, and postictal signal traces, meaning that there is an imbalanced class representation for the binary and multi-class classification problems. Consequently, we implement a data balancing algorithm that guarantees there is a balanced representation of classes. For this, we consider the maximum number of ictal samples within a run to be the total number of ictal samples, and non-ictal samples are taken from the preictal and postictal traces in equal amounts. This process is illustrated in Figure 5. Each data sample represents a GR that is used for the train, validation, and test datasets, marked with a red cross, a cyan square, and a turquoise circle, respectively.

**Figure 5.**
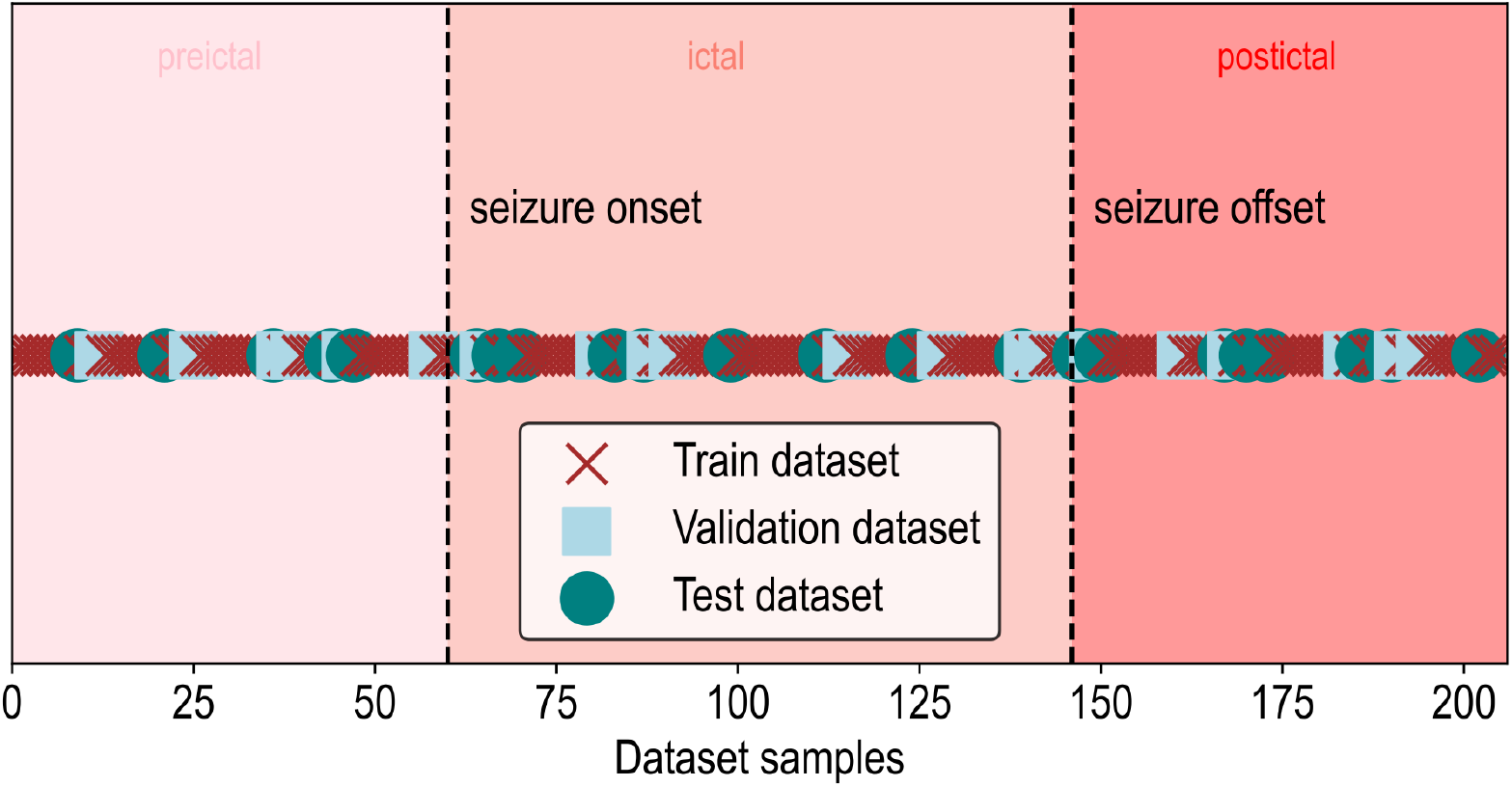
Illustration of the data balancing procedure.

#### 2.2.6 Graph neural network model

The GNN model proposed by Grattarola et. at. in [8] is illustrated in Figure 6. This depicts a 2-layer neural network composed of an ECC layer [13] followed by a GAT layer [14]. The ECC layer constrains the learning to focus on the edge features, while the GAT layer computes attention coefficients, which are measurements that quantify the importance of each node in a graph in the learning process. In [8], the authors proposed the use of the attention mechanism as a means to map attention coefficients to the SOZ. However, we did not find success in applying this methodology, and we plan to devise strategies for SOZ localization within our pipeline in future work.

**Figure 6.**
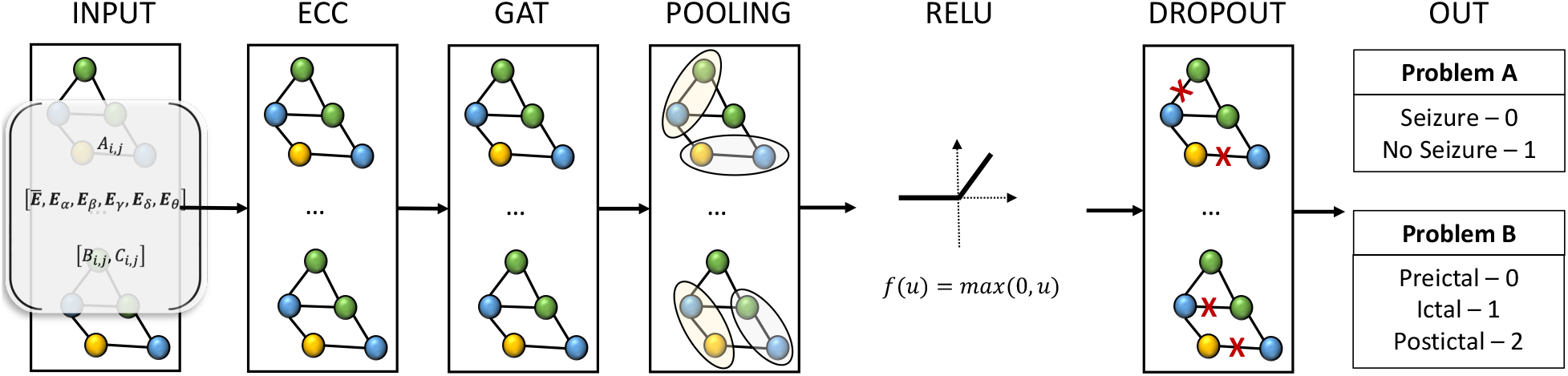
Architecture of the GNN model proposed by Grattarola et. at. in[8].

## 3 Results

### 3.1 Binary classification of ictal and non-ictal activity

In Figure 7 we show the performance evaluation of the GNN model using the 9 GR tests introduced in Table 2. We consider 4 metrics to assess the performance of the GNN model: accuracy, F1-score, AUC, and loss, shown in Figures 7a, 7c, 7b, 7d. We can clearly see that by using the proposed features from the iEEG data into the GRs we can significantly improve the performance of the GNN model with accuracy, F1-score, and AUC with a median near 90% for tests t21, t22, and t23. Also, we can see that the performance becomes more consistent as more features are considered.

**Figure 7.**
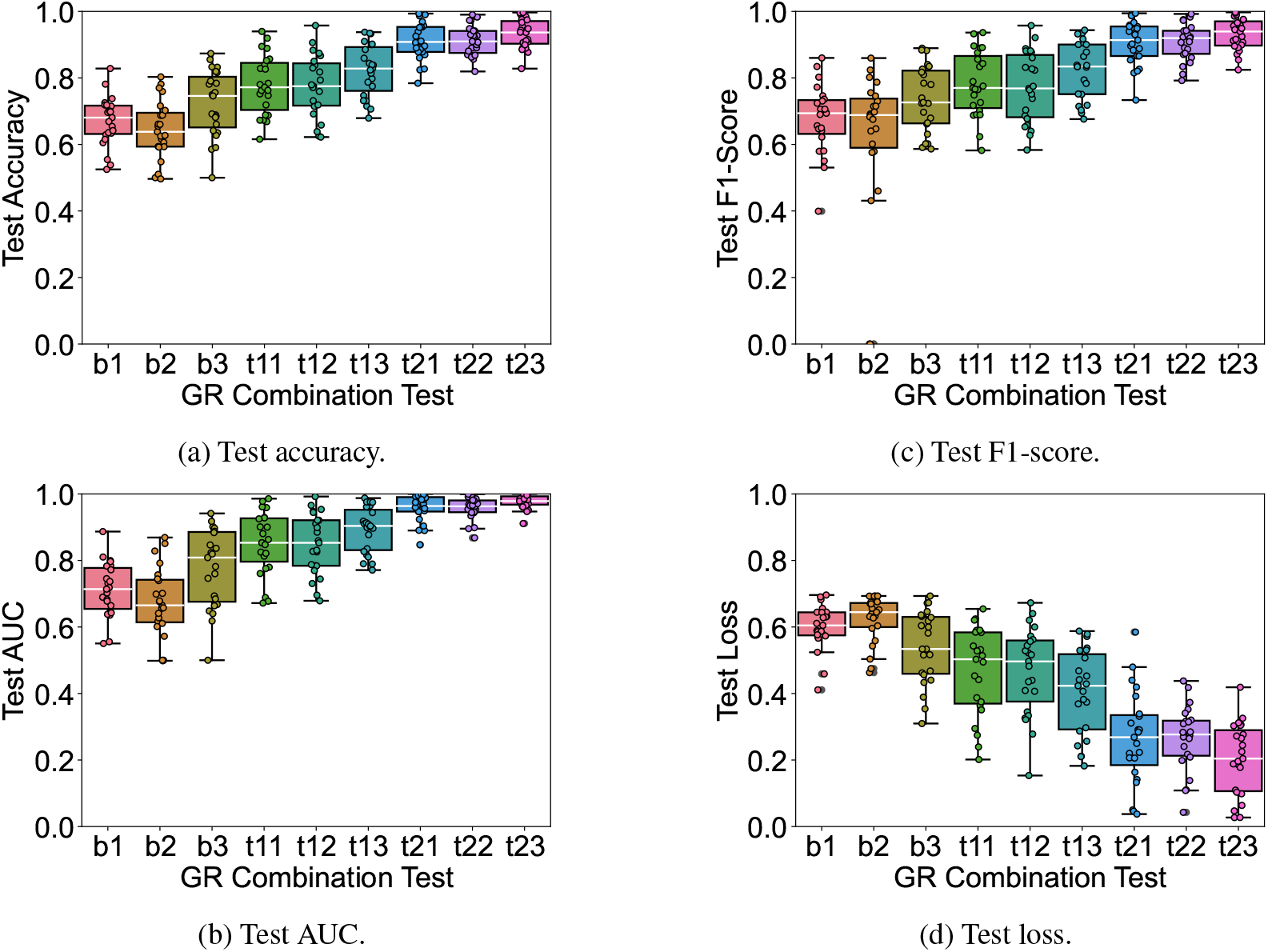
Test results performance for the binary classification seizure detection problem.

However, our methodology is limited to the features that we use for GR creation (shown in Table 2). Thus, further work is needed to evaluate other FCN-based features that may inform of higher-level processes, such as the direct absolute coherence (DAC) [30] that adds directionality of signaling to the network abstraction, which we plan to investigate in future work.

### 3.2 Multi-class classification of preictal, ictal, and postictal activity

In Figure 8 we show the evaluation of the GNN model using the same GRs from the binary classification problem for the multi-class classification problem. Here, we also see that by including more data within the GRs we can significantly improve the performance of the GNN model. Importantly, we can see that the GNN model is capable of discriminating between preictal and postictal signal traces, showing a potential to develop seizure prediction algorithms in the future. However, it is not yet clear which GR is more suitable for every patient iEEG record.

**Figure 8.**
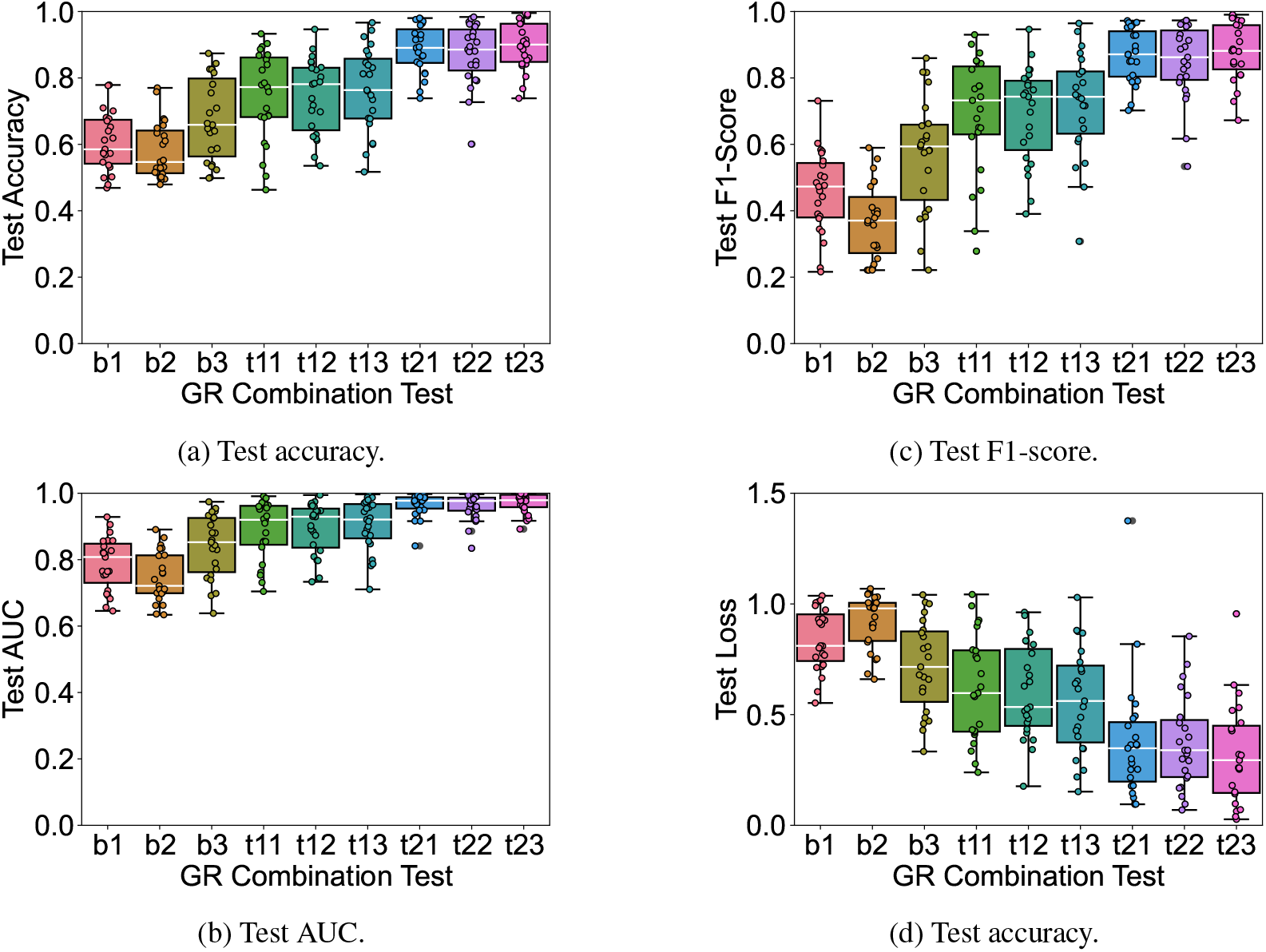
Test results performance for the multi-class classification seizure detection problem.

In general, our methodology is better than the baseline, however, same GR structures do not work equally well for all patients, as depicted by the outliers in this figure. We speculate that similarly to the difficulties encountered by clinical experts in visually inspecting the iEEG data, GNN models struggle to learn features from ambiguous or corrupted data recordings. In future work, we will investigate the relationships between graph embeddings and iEEG signal characteristics, and their impact on GNN learning and decision-making processes.

## 4 Concluding remarks

We have shown an iEEG processing pipeline that creates GRs of iEEG data, and we used them to build a GNN model that can detect ictal and nonictal activity, as well as distinguishing between preictal, ictal, and postictal activity. In this section, we want to discuss some of the challenges that we faced in dealing with the data and automating these processes, as well as to suggest future steps to continue with this work.

### 4.1 On the data

Accessing the data was hard, and from the reported 100 patient data available online, we were only able to use 25 patient records. A large portion of the data was simply not available because one of the research centres failed to de-identify and share. In the OpenNeuro repository there are 35 data entries. However, for the purposes of our study, 10 entries were not usable due to inconsistencies in the clinical annotations. For example, in some cases there were not marks for the seizure onset or the seizure offset events.

An interesting observation that we noted in the data was with regard to the clinical annotations of the seizure onset and seizure offset marks. For clinical purposes, identifying the beginning and the end of a seizure consists in marking the moment in time when a seizure starts at few electrodes, resulting in clinical annotations as shown in Figure 1. However, it can take several seconds for a seizure to spread to multiple regions. While this does not hinder the ability of clinical experts to make decisions based on the data and their annotations, we consider this to be a major problem for building systems to automate the process of seizure detection. The reason being that the data at its core is mislabeled. While the clinical annotation considers one general time point as the onset of a seizure event, the spread of the seizures to other brain regions as seen by the iEEG electrodes is registered several seconds after the clinical mark. Consequently, our results are currently affected by this label noise. As future steps, we will work with EEG technologists at the Toronto Western Hospital to devise a seizure detection algorithm to re-label clinical data and make up for these inconsistencies, which we believe will improve the performance and reliability of the current GNN model.

### 4.2 On the functional connectivity measurements

The three different methods that we consider for computing FCNs of iEEG data focus on different abstractions of the data. For instance, the Pearson correlation allows us to quantify similarities in signal activity given by the energy levels at any point in time, which is a useful abstraction to study iEEG patterns from an energy perspective in the time domain. Similarly, the coherence also allows us to quantify energy levels but in the frequency domain, thus allowing us to study iEEG patterns in the frequency domain. And, the PLV allows us to quantify synchronicity across signals over time. In this paper, we have shown that these functional connectivity measurements are strong representations of the iEEG data so that artificial neural networks can *learn* from the general dynamics of the data. The next step that we will study is what exactly are the networks learning from the data, to determine how to trace it back to the SOZ.

### 4.3 On the graph representations

We only considered 9 GRs with 3 different adjacency matrices, 2 different node and edge features, and tested them for 2 classification problems. Thus, further work will be required to determine what is the most powerful GR of iEEG data to solve the problems of seizure detection and SOZ localization. In addition, further work will be required to determine what other uses of GRs of iEEG data can be found for assisting the treatment of other neurological disorders such as Parkinson’s disease, Alzheimer’s disease, or major depression disorder.

### 4.4 On the graph neural networks model

The field of GNNs has been growing at a fast pace in recent years. Thus, further work will be required for the evaluation of more GNN architectures to unlock their full potential for solving the problems of seizure detection and SOZ localization, and for their application to neuroscience in general.

Moreover, we are developing self-supervised learning (SSL) processes within our pipeline. SSL is a paradigm of deep learning that has proven excellent at uncovering patterns from structured data in the fields of computer vision and large language models [31–33]. Recently, few works have shown the potential of SSL to uncover patterns from scalp EEG data[34].

Automating the process of iEEG-based seizure detection within clinical pipelines treating epilepsy is a timely need to improve healthcare practice. In this paper we proposed an iEEG data processing pipeline that uses a GNN model to detect ictal and non-ictal activity, and that detects preictal, ictal, and postictal data. Moreover, we demonstrate that by leveraging iEEG signal data as GRs of iEEG data we can significantly improve the performance of GNN-based systems.

## Acknowledgements

We thank Professors Taufik Valiante and Mary Pat McAndrews at the Krembil Research Institute for their valuable input and support in the conceptualization of this project. As well, we thank Dr. Daniele Grattarola for sharing his technical expertise in the development of graph neural network models.

## Notes

### Competing Interest Statement

The authors have declared no competing interest.

